# Bridging psychology and genetics using large-scale spatial analysis of neuroimaging and neurogenetic data

**DOI:** 10.1101/012310

**Authors:** Andrew S. Fox, Luke J. Chang, Krzysztof J. Gorgolewski, Tal Yarkoni

## Abstract

Understanding how microscopic molecules give rise to complex cognitive processes is a major goal of the biological sciences. The countless hypothetical molecule-cognition relationships necessitate discovery-based techniques to guide scientists toward the most productive lines of investigation. To this end, we present a novel discovery tool that uses spatial patterns of neural gene expression from the Allen Brain Institute (ABI) and large-scale functional neuroimaging meta-analyses from the Neurosynth framework to bridge neurogenetic and neuroimaging data. We quantified the spatial similarity between over 20,000 genes from the ABI and 48 psychological topics derived from lexical analysis of neuroimaging articles, producing a comprehensive set of gene/cognition mappings that we term the Neurosynth-gene atlas. We demonstrate the ability to independently replicate known gene/cognition associations (e.g., between dopamine and reward), and subsequently use it to identify a range of novel associations between individual molecules or genes and complex psychological phenomena such as reward, memory and emotion. Our results complement existing discovery-based methods such as GWAS, and provide a novel means of generating hypotheses about the neurogenetic substrates of complex cognitive functions.

## Introduction

It is widely held that thoughts, feelings, and actions are reflected in macroscopic neural patterns that emerge from microscopic molecular processes that orchestrate the function of our nervous system. Although the basic concept of a matter-based mind is no longer a matter of serious scientific debate, the precise mapping between molecules and mental states remains largely a mystery. A major barrier to progress is that psychological and molecular processes unfold on vastly different spatial and temporal scales. In fact, the chasm between the two levels of description may be too wide to bridge directly, and has resulted in the emergence of multiple non-overlapping scientific fields. In the present work, we demonstrate the utility of distributed macroscopic neural patterns as a novel means of bridging the long-standing gap between psychological and molecular neurobiological levels of analysis.

Brain-wide macroscopic spatial patterns are ideal for linking cognitive and molecular processes because of their accessibility to multiple disciplines and levels of analysis (*1–3*). In recent decades, functional neuroimaging studies have identified highly consistent distributed brain networks that underlie mental states ranging from reward-seeking (*4*) to goal-directed thought (*5*) to autobiographical memory (*6*). Simultaneously, animal and human studies involving positron emission tomography, autoradiography, and in situ hybridization have demonstrated that many genes and the proteins they code for have predictable large-scale patterns of expression throughout the human brain (*7–10*). For example, in support of the role of dopamine in reward processing, it has been observed that genes encoding dopamine receptor proteins implicated in reward processing are highly expressed in consistent areas of the mammalian striatum(*11, 12*). These regions are reliably activated in fMRI studies of reward processing (*13, 14*), and show extremely high concentrations of dopamine DRD2/DRD3 receptors in *in vivo* human imaging studies using positron emission tomography (PET)(*15*). The convergence of psychological and molecular processes at the level of large-scale, brain-wide spatial expression thus offers a powerful potential window into the molecular bases of cognition and affect.

In the work reported here, we introduce, validate, and apply a novel tool for mapping cognitive phenomena to molecular processes based on large-scale spatial analysis. We harness state-of-the-art neuroimaging and neurogenetics databases: Neurosynth, our recent framework for large-scale, automated synthesis of the published fMRI literature (*16*), and the Allen Human Brain Atlas (AHBA)(*17*), a brain-wide gene expression atlas derived from transcriptome-wide mroarray assessments of human brain tissue. Here, we use this spatial integration of molecular and psychological processes (Neurosynth-gene) approach to independently replicate previous associations identified in the experimental literature, and to identify a large number of novel associations that have not been previously reported. This non-mechanistic approach can help guide researchers toward the most productive avenues for future research aiming to understand how complex psychological phenomena such as reward and memory processing emerge from the microscopic molecules of the brain.

## Results

### A common space for gene expression and functional activation

Our work builds directly on the Neurosynth framework (*16, 18, 19*). Neurosynth is an open framework for automated synthesis of published fMRI results, and enables researchers to produce high-quality estimates of the brain-wide neural correlates of major cognitive tasks and psychological states. The framework is ideal for understanding the relationship between psychological constructs and the brain, as it provides quantitative inferences about the consistency and specificity (*20, 21*) with which different cognitive processes elicit regional changes in brain activity. It can, for example, generate maps that estimate the relative likelihood with which activation in a given brain region implies the presence of a particular psychological process such as reward or emotion, enabling “decoding” of entirely novel images in a relatively open-ended way (Figure 1a)(*18, 22*). Currently, Neurosynth provides whole-brain maps for several thousand distinct terms; in the present analyses we used a dimensionally-reduced set of 48 topics that reflect high-level psychological constructs such as Emotion, Memory and Reward (Fig 1b; Supplementary Table 1; Supplementary Figure 1; Supplementary Methods).

**Figure 1.**
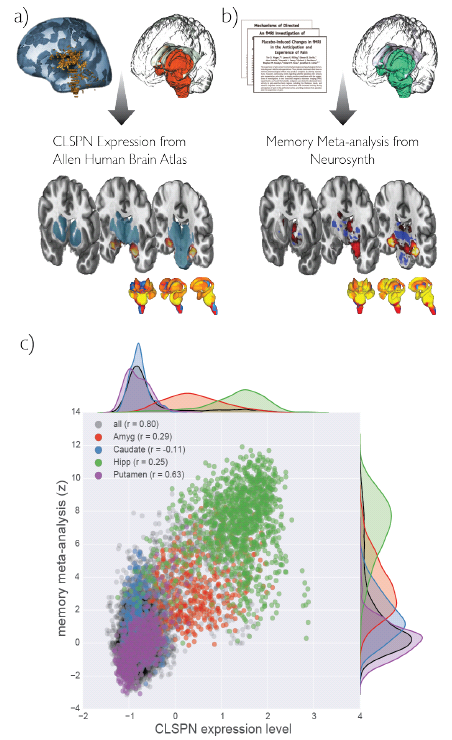
The Neurosynth-gene atlas uses spatial associations to link genes to cognitive processes, and is exemplified here by CLSPN gene expression from the AHBA and Memory meta-analysis map in Neurosynth. (a) Gene expression levels were extracted from the AHBA for subcortical structures and projected into standard space, shown here for the gene CLSPN. (b) Text mining is used to automatically generate “reverse inference” meta-analytic maps of fMRI studies within the same subcortical regions, exemplified here for the Memory topic. Patterns of gene expression and meta-analytic statistics are shown on both 2d slices and 3d renderings of subcortical structures (c) Scatter and kernel density estimation plots displaying spatial relationship between CLSPN expression levels and memory-related activation across all subcortical voxels. Different brain regions are represented in different colors.

To bridge between large-scale functional activation and underlying molecular mechanisms, we integrated Neurosynth with data from the recently-released AHBA (*23*). The AHBA is a brain-wide gene expression atlas derived from transcriptome-wide microarray assessments of brain tissue from 3702 samples collected across 6 human donors. It provides a window into the distribution of human gene expression throughout the adult brain—an ability that has already led to novel insights (*24*). Gene expression is ideal for large-scale examinations of the molecular composition of neural tissue, as transcription of genes into RNA is a critical step in converting each cell’ s DNA into the proteins that determine its function. Gene expression depends on genetic structure, the local molecular environment, and epigenetic factors, and, although each cell contains a full genome, patterns of gene expression are specific to particular cell types. Local regulation of gene expression is critical for determining the structure and function of neurons and glia by altering the composition of the cell, and thus gives rise to brain-region specific functions.

To quantify the spatial similarity between gene expression and functional activation maps, we transformed the gene expression data from the AHBA into the common stereotactic brain space used by Neurosynth. This process consisted of (i) normalizing brain-wide expression values separately for each gene averaged across probe-sets, (ii) mapping the reported coordinates for brain tissue used for microarray analyses to a standard neuroimaging template-space (i.e. Montreal Neurological Institute), and (iii) smoothing the data to match the resolution of the Neurosynth maps (see Supplementary Methods for details). Because gene expression patterns differ substantially between subcortex and cortex (*17*), we conducted separate subcortical and cortical analyses. Here we focus exclusively on sub-cortical inferences, as the current spatial distribution of the AHBA samples was sparse in cortex (see Supplementary Methods; Supplementary Figure 2). Once the AHBA and Neurosynth maps were represented within the same standard brain space, we computed the spatial correlation between each gene expression map and each psychological topic map. The resulting matrix of 29,180 × 48 associations (i.e., genes × topics) – which we term the Neurosynth-gene atlas–provided a rich substrate for subsequent hypothesis testing and exploration of the relationship between specific genes or gene families and broad cognitive and affective processes.

**Figure 2.**
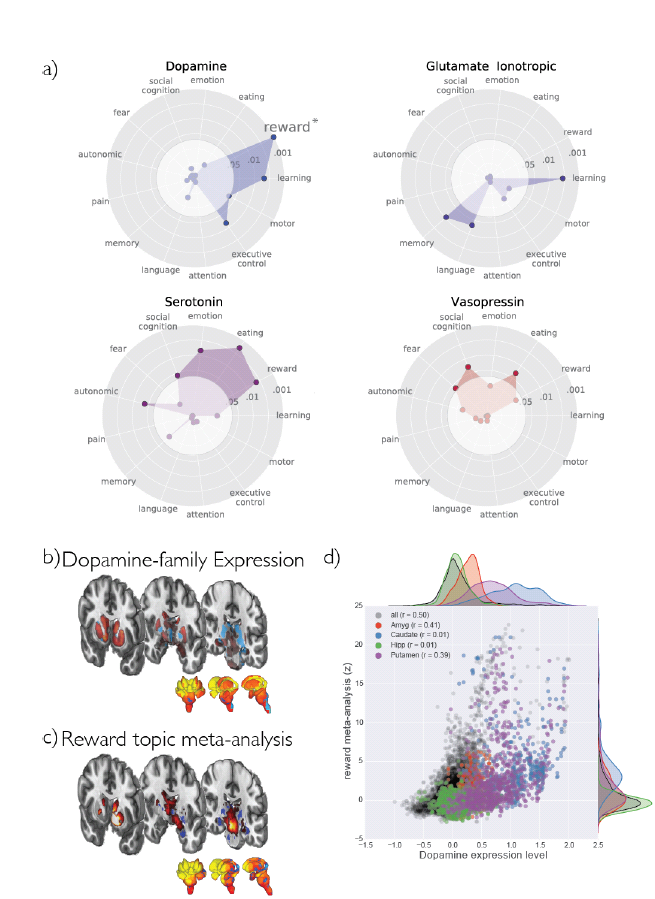
Neurotransmitter-genes are selectively associated with Neurosynth topics. Radar plots displaying statistical association between selected HGNC neurotransmitter receptor gene groups and selected Neurosynth topics (a) (for full results, see Supplementary Figure 4). Associations in the dark outer circle are significant at p < .05. Each statistical association represents the likelihood, relative to other neurotransmitters, that groups of neurotransmitter receptor genes are expressed in topic-related areas. For example, the dopamine-reward relationship (denoted with an asterisk) reflects the fact that patterns of gene expression in the Dopamine gene family (b) and the Reward-topic meta-analysis (c) are correlated across the brain (d).

To illustrate, Figure 1 displays the single strongest spatial correlation in the entire database—between expression of the gene CLSPN (claspin) and Memory-related brain activation—(r = .80). Importantly, this correlation did not solely recapitulate anatomical boundaries (i.e., that CLSPN simply happens to be more strongly expressed in the hippocampus, which is known to be implicated in memory formation), as the positive correlation between patterns of CLSPN expression and memory-related activation is apparent both within and across multiple brain structures (Fig. 1C).

### Strong corroboration of consensus neurotransmitter-cognition associations

The ability to search for candidate genes by psychological constructs (or for candidate psychological constructs by gene) presents a powerful tool for mapping between molecular and cognitive levels of analysis. The Neurosynth-gene atlas can be used for both top-down, theory-driven testing of hypotheses regarding the distribution of gene expression in the brain regions implicated in a specific cognitive process, as well as bottom-up, data-driven hypothesis generation aimed at identifying novel gene-cognition relationships.

To demonstrate the utility of the Neurosynth-gene atlas, we first sought to corroborate existing relationships between neurotransmitter systems and psychological processes—e.g., the well-established link between dopamine and reward (*25*), and the heavily-studied, though more contentious, relationship between the serotonin system and depression (*26–28*). Neurotransmitter systems are an ideal target for this kind of validation, as numerous studies have examined the psychological effects of specific pharmacological neurotransmitter receptor activation. Table 1 lists 16 frequently studied (though certainly not exhaustive) neurotransmitter/cognition associations--many of which have been reported in thousands of studies (e.g., dopamine/reward, serotonin/depression, and oxytocin/social behavior). To assess the relative strength of each of these associations, we first selected all individual genes contained in the corresponding HUGO Gene Nomenclature Committee (HGNC) neurotransmitter receptor group (e.g., DRD1, DRD2, etc.; see Supplementary Methods). We then computed the mean correlation with the target Neurosynth topic for all genes within each neurotransmitter group, and used a permutation-based statistical approach to quantify the likelihood of obtaining an association of the observed strength purely by chance (see Supplementary Methods). The results supported 12 of the 16 hypothesized relationships (Table 1), effectively “re-discovering” these known associations using the Neurosynth-gene atlas.

**Table 1.** The Neurosynth-gene atlas was able to identify 12/16 frequently studied neurotransmitter-cognition associations. This non-exhaustive list was derived from PubMed abstracts related to neurotransmitters and cognitive science topics, verified and collated by ASF, LJC & TY. R-values indicate the mean partial correlation between genes in the gene-receptor family (e.g. Dopamine) in relation to the Neurosynth topic (e.g. reward) across the thousands of subcortical voxels. P-values were computed using a permutation analysis (see supplementary methods for details).

| # | Neurosynth-gene atlas<br>found<br>support | Receptor<br>type | Putative<br>cognitive<br>function | Neurosynth<br>topic map | mean<br>r-value<br>across<br>subcorte<br>x | p |
| --- | --- | --- | --- | --- | --- | --- |
| 1 | yes | Dopamine | reward | reward | 0.23 | **0.0004** |
| 2 | yes | Dopamine | working<br>memory | working<br>memory | 0.08 | **0.03** |
| 3 | no | Dopamine | motor<br>function | motor<br>processing | 0.08 | 0.08 |
| 4 | yes | Dopamine | ADHD | ADHD | 0.14 | **0.002** |
| 5 | yes | Serotonin | depression | depression | 0.07 | **0.002** |
| 6 | yes | Serotonin | emotion | emotion | 0.05 | **0.02** |
| 7 |  | Opioids | pain | pain | 0.06 | 0.34 |
| 8 | yes | Vasopressin/<br>Oxytocin | social<br>behavior | social<br>processing | 0.1 | **0.03** |
| 9 | no | Acetylcholine<br>(Muscarinic) | learning<br>and<br>memory | learning | 0.01 | 0.33 |
|  |  |  |  | memory | -0.07 | 0.69 |
| 10 | yes | Acetylcholine<br>(Nicotinic) | working<br>memory | working<br>memory | 0.05 | **0.02** |
| 11 | yes | Acetylcholine<br>(Nicotinic) | attention | attention | 0.06 | **0.02** |
| 12 | yes | Glutamate<br>(Ionotropic) | learning<br>and<br>memory | learning | 0.11 | **0.006** |
| 13 | yes | Glutamate<br>(Ionotropic) |  | memory | 0.15 | **0.0005** |
| 14 | yes | Neuropeptide Y | feeding | feeding | 0.11 | **0.005** |
| 15 | yes | Neuropeptide Y | Anxiety/<br>Fear | anxiety | 0.11 | **0.02** |
|  |  |  |  | fear | 0.01 | 0.46 |
| 16 | no | GABA | Anxiety/<br>Fear | anxiety | -0.03 | 0.95 |
|  |  |  |  | fear | -0.03 | 0.95 |

To assess the specificity of these associations and ensure that these positive results did not reflect broader relationships (e.g., that dopamine was spatially correlated with several cognitive processes because the latter were all themselves highly intercorrelated), we expanded our analysis to include all possible pairs of HGNC neurotransmitter receptor groups (37 in all; Supplementary Figure 3) and Neurosynth topics. Most of the confirmed associations reported in Table 1 displayed a striking degree of specificity in this latter analysis (Figure 2). For example, while we had expected that dopamine receptor genes would be preferentially expressed in subcortical brain regions associated with reward, we did not necessarily expect the association to be highly selective, and had anticipated that a number of other neurotransmitters would also show strong associations with reward. Yet of 37 distinct neurotransmitter receptor families,

Reward was most strongly associated with the DRD family (p = 0.0004), and the only other significantly associated neurotransmitter group was serotonin (a neurotransmitter also implicated in reward; p = 0.02; Fig. 2; all neurotransmitter/topic relationships can be seen in Supplemental Figures 4 and 5). Examination of the spatial intercorrelations between Neurosynth topic maps further demonstrated low correlations between most maps (Supplementary Figure 6)–for example, the spatial intercorrelations between the Reward, Working Memory, and ADHD maps did not exceed 0.16, even though the DRD family was significantly associated with all three topics. Thus, these results validate the use of spatial expression mapping as a bridge between molecular genetics and human cognition by providing strong independent replications of associations that have previously been demonstrated using very different methodological approaches.

### A discovery tool for gene-family-cognition associations

The successful replication of previously established association carries with it an important implication: if the present approach can successfully recapture known relationships, it is likely to also have considerable utility in testing other, more speculative, hypotheses, as well as in identifying entirely novel associations. Consistent with this notion, we observed a number of statistically reliable gene-cognition associations that, to our knowledge have not been previously reported, and yet are broadly consistent with existing literature (as seen in Supplementary Figure 5). For example, ‘Pain’ was associated with the Sphingosine gene family, members of which have been suggested to play a role in pain signaling (*29*), and ‘Social Cognition’ was associated with the Trace Amine gene family, members of which have been linked to the processing of socially-relevant smells (*30*).

Next, we generalized our approach beyond neurotransmitter genes by conducting a comprehensive analysis of 397 gene families labeled in the HGNC. The results confirmed the striking selectivity of many of the neurotransmitter effects reported above. For example, of 397 gene groups, the single strongest spatial similarity to the Neurosynth Reward map was observed for the dopamine receptor family (i.e. DRD; p < .0001), and the single strongest similarity to the Emotion map was observed for the ionotropic serotonin receptor family (HTR3; p < .0001). Multiple comparison correction revealed other associations previously reported in the literature (Supplementary Figure 7). For example, the Neurosynth Memory map was significantly associated (*p* < .001) with the pattern of expression for genes that code for constituent proteins of the Actin-related protein 2/3 complex (Arp2/3), consistent with recent reports implicating the Arp2/3 complex in memory formation and forgetting (*31–34*).

### Hierarchical clustering of gene/cognition associations

The preceding family-based analyses all assume strong prior knowledge about the grouping structure of individual genes. However, such a top-down approach risks overlooking any structure in the covariance of gene/cognition correlations that does not respect the boundaries of existing gene ontologies. For example, even within the DRD gene group—which displayed strong relationships with reward in the aggregate—there is known heterogeneity: the D4 dopamine receptor is known to show an affinity for other catecholamines besides dopamine (*35, 36*). A major advantage of a discovery tool like Neurosynth-gene is its potential to complement and inform existing ontologies by deriving novel, data-driven, gene clusters. To this end, we used hierarchical clustering to identify groups of neurotransmitter receptor genes that exhibited similar profiles of association with Neurosynth topic maps irrespective of their nominal family membership in the HGNC ontology (Figure 3; full clustering results can be seen in Supplementary Figure 8). These analyses revealed that most, but not all, dopamine receptors concentrated together within a single cluster that loaded strongly on the Reward and Learning maps. Interestingly, DRD4 was not included in this cluster; instead, it fell into a cluster of genes that loaded on the Attention and Poly-modal Sensory topics. Moreover, these analyses provide insight into other, less-well understood neurotransmitters. For example, some (P2RY1 and P2RY11), but not all (P2RY12 and P2RY13), of the P2Y purinoceptors were concentrated within a Reward-related cluster, which is both consistent with the proposed role of P2Y in addiction, as well as the known heterogeneity within this system and the molecules bound by these receptors (*37*).

**Figure 3.**
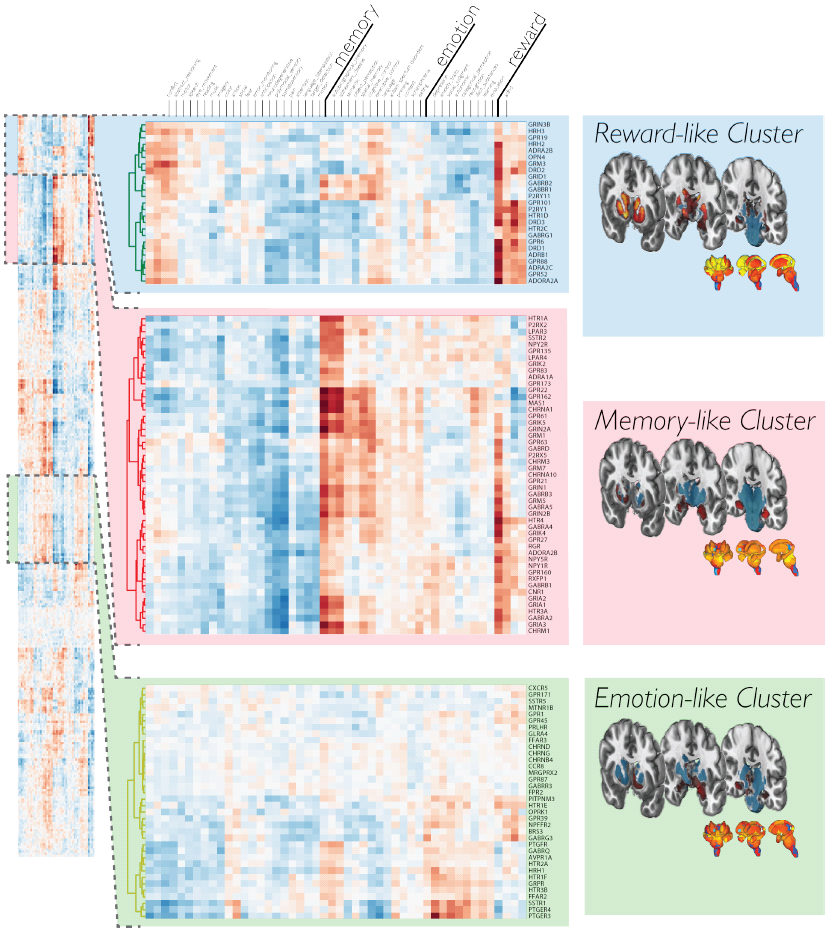
Hierarchical clustering of neurotransmitter receptor genes based on spatial similarity to Neurosynth topics, revealed clusters that span neurotransmitter groups. Clustergram for all 315 genes (left), with an expanded view displaying three example clusters (middle) that load highly on reward, memory, and emotion, respectively. Subcortical renderings of gene expression levels, averaged over all genes within each cluster can be seen on the right. The full clustergram can be seen in Supplementary Figure 8.

### A discovery tool for individual gene-cognition associations

Finally, we turned to what is arguably the most tantalizing use of our gene-cognition mapping approach: the potential to conduct data-driven Neurosyth-gene atlas searches for associations between individual genes and specific cognitive or affective processes. One way to conduct such an exploration is to inspect the genes most strongly correlated with target cognitive and affective processes. Figure 4 displays the 10 individual genes most strongly associated with selected Neurosynth topics (for additional results, see Supplementary Figure 9). Not surprisingly, many of the gene-level findings recapitulated the family-level results (e.g. DRD3, seen in Supplementary Figures 9 and 10). For example, consistent with the family-level correlations for the DRD gene group, we identified a strong correlation across subcortex between DRD3 expression and Reward (r = 0.55). In fact, DRD3 expression was more strongly associated with Reward than all but two other genes (GUCA1A and GPR101). A similarly strong association (r = .58) was observed between learning and the A2A adenosine receptor gene, which animal models have implicated in multiple forms of learning and habit formation (Supplementary Figure 10) (*38–40*).

**Figure 4.**
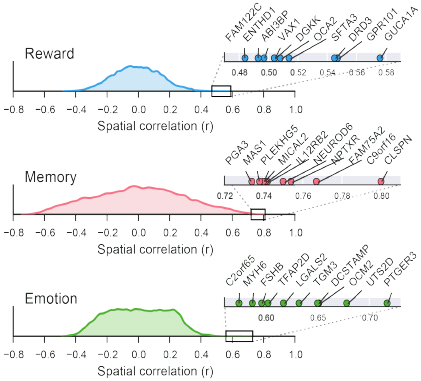
Novel gene-cognition association discovery. Kernel density estimation plots displaying the distribution of spatial correlations with all genes for three sample Neurosynth topics, Reward, Memory and Emotion. Right: zoomed-in view of the top 10 individual gene correlations for each topic.

Importantly, in addition to these examples, we observed numerous other gene-level associations that were not subsumed by the family-level analysis yet converged with prior theoretical and empirical work (for this and other example relationships depicted in scatter plots see: Supplementary Figure 10). Notable associations include Fear and SSTR1 (Somatostatin Receptor Type 1), which has been implicated in the genetics of panic disorder, used to treat patients with panic, and implicated in animal models of fear (*41–43*); pain and LGALS1, a Galectin-1-coding gene implicated in the development of acute and chronic inflammation in knockout models (44–46); and motor control and several genes located within the spinal muscular atrophy gene region at 5q13.1 —including SMN1 (Survival of Motor Neuron 1), the putative locus of causal effect (*47–49*).

Interestingly, we also identified a large number of gene-cognition relationships which are not obviously supported by the extant literature. These novel relationships are particularly exciting because they provide an impetus to investigate novel associations between psychological constructs and their molecular bases that might otherwise go unstudied. For example, if future research demonstrates a causal role for the aforementioned claspin-memory association (Figure 1), this would provide an unpredicted link between claspin-dependent processes and the formation of new memories. One might speculate, for instance, that the known role of claspin in the maintenance of genome integrity (*50, 51*) plays a critical role in the high-fidelity genome duplication that is required for memory-related hippocampal neurogenesis (*52, 53*).

### Discussion

Understanding the molecular basis of human cognitive processes promises to illuminate the biology that embodies our thoughts and emotions. In this regard, identifying the genes that alter expression of the proteins that comprise cells and synapses—giving rise to the patterns of brain activation that underlie complex cognitive phenomena—is critical. Here we demonstrated that patterns of spatial covariation between gene expression and meta-analytic functional brain activity can provide a unique and powerful window into the relationships between genes and cognition. Our findings independently replicate numerous prior gene-cognition associations, and provide a novel discovery tool for identifying molecules that may participate in specific psychological or cognitive processes. This approach complements other discovery-based methods such as GWAS, and aims to accelerate future mechanistic research by generating novel hypotheses.

Our contention is not, of course, that all—or even most—such associations are likely to accurately reflect a role of specific gene products in human cognition—but rather, that an as-yet undetermined subset of them undoubtedly do. Moreover, as more data become available, the present findings will improve in tandem. While the Allen Human Brain Atlas is a remarkable resource, it is important to remember that it presently contains relatively sparsely sampled data from only six human brains (particularly in cortex as seen in Fig S2c). As the density and quality of the AHBA dataset increases, we anticipate that the sensitivity and specificity of the gene-cognition mappings reported here will also improve—potentially dramatically. Moreover, datasets documenting gene expression changes as a function of age, individual variation, and/or context will only increase the potential for the Neurosynth-Gene approach. In the meantime, to facilitate further development and application of our methods, we have made all of the software, data, and results used to produce these findings publicly available on the web (http://github.com/neurosynth/neurosynth-genes). We also provide interactive, downloadable, whole-brain gene expression and functional activation maps via our Neurosynth web interface (http://neurosynth.org; Supplementary Figure 11). Our hope is that geneticists and cognitive neuroscientists will use these new resources for both theory-driven hypothesis testing and bottom-up discovery of a wide range of gene-cognition associations.

## Acknowledgements

We would like to thank L. Ruzic, Y Ashar, W. Moore, N.J. Vack, P. Roseboom, J. Oler, and D. Margulies for their helpful consideration of this approach. This work was funded by NIH R01MH096906.

